# Cryo-EM of a *Marseilleviridae* virus particle reveals a large internal microassembly

**DOI:** 10.1101/139097

**Authors:** Kenta Okamoto, Naoyuki Miyazaki, Hemanth K.N. Reddy, Max F. Hantke, Filipe R.N.C. Maia, Daniel S. D. Larsson, Chantal Abergel, Jean-Michel Claverie, Janos Hajdu, Kazuyoshi Murata, Martin Svenda

## Abstract

Nucleocytoplasmic large DNA viruses (NCLDVs) blur the line between viruses and cells. Melbournevirus (MelV, fam. *Marseilleviridae*) belongs to a new family of NCLDVs. Here we present an electron cryo-microscopy structure of the MelV particle, with the largest known triangulation number (T=309) for a virus. The 230-nm particle is constructed by 3080 pseudo-hexagonal capsomers and encloses a membrane bilayer. Its most distinct feature is a large dense body (LDB) consistently found in all particles. Electron cryo-tomography of 147 particles showed that the LDB is located preferentially in proximity to the bilayer. The LDB is 30 nm in size and its density matches that of a genome/protein complex. More than 58 proteins are associated with the purified particle, including histone-like proteins, putative membrane proteins and capsid proteins. The observed intricate structural organization reinforces the genetic complexity of MelV, setting it apart from other viruses, and suggests an evolutionary link with cellular organisms.

## Introduction

Nucleocytoplasmic large DNA viruses (NCLDVs) share genetic and structural traits, and propagate widely in free-living unicellular microorganisms such as protozoa and some algae (Iyer et al., 2001). Comparative genomics of NCLDVs has evoked speculations on the origin of DNA viruses as a distinct domain of life and their role in the evolution of cellular organisms (Abergel et al., 2015, Claverie and Abergel, 2013).

The initial representatives of the icosahedral NCLDVs infecting unicellular eukaryotes were isolated from Chlorella, a fresh water algae. These include the 190 nm in diameter *Paramecium bursaria* chlorella virus type 1 (PBCV-1), the 220 nm in diameter *Phaeocystis pouchetii* virus (PpV01) and the 150 nm in diameter *Phaeocytis globosa* virus (PgV), respectively (Van Etten et al., 1982, Monier et al., 2008, Yan et al., 2005, Santini et al., 2013). Fourteen years ago, the much larger amoebal virus, Mimivirus, was identified from the water in a cooling tower of a hospital in England (La Scola et al., 2003). The Mimivirus particle forms a pseudo-icosahedral core of 450 nm in diameter covered with long fibers, 150 nm in length (La Scola et al., 2003, Xiao et al., 2005). Since the discovery, a vast number of Mimivirus-like viruses have been isolated from fresh water, seawater, and soil samples, as well as from living organisms (Claverie et al., 2009, Saadi et al., 2013, Dornas et al., 2014). *Marseilleviridae* is a recently established family among the large amoebal DNA viruses (Colson et al., 2013). They possess capsid structures that vary in size from 190 to 250 nm. The particle architecture and the genome complexity classify them into a unique family separated from the *Mimiviridae* viruses (Aherfi et al., 2014b, Doutre et al., 2014, Doutre et al., 2015, Thomas et al., 2011). Faustovirus, another recently isolated large amoebal virus, is classified into yet another new family of the NCLDVs. It is phylogenetically close to African swine fever virus (ASFV) (Reteno et al., 2015). All these NCLDVs seem to share an evolutionary relationship, albeit complex, based on a core set of conserved genes and their virion structures (Iyer et al., 2001, Koonin and Yutin, 2010, Yutin and Koonin, 2012). Their genomes encode a conserved major capsid protein (MCP). The MCPs form pseudo-hexagonal trimers of approximately 70 Å in diameter, which is the constructing unit (capsomer) in the capsid lattice (Nandhagopal et al., 2002).

Structural studies of the NCLDV particles are important for understanding their assembly, mechanism of cell entry and evolution, but the challenges lie in their large sizes. Few of the large algal and amoebal virions have been structurally analyzed. The first electron cryo-microscopy (cryo-EM) structure of NCLDVs revealed that the large capsid adopts an icosahedral lattice with a large triangulation number (T-number). The pseudo-hexagonal capsomers form pentasymmetron and trisymmetron superstructures (Yan et al., 2000). The capsid structures of the algal PBCV-1 and PpV01 with T=169 and T=219 symmetries were determined to around 30 Å resolution using cryo-EM single particle analysis (SPA) (Yan et al., 2005, Yan et al., 2000). Recently, the two capsid layers of the amoebal Faustovirus with T=64 (inner layer) and T=277 (outer layer) symmetries were determined to 15 and 26 Å resolution, respectively (Klose et al., 2016). The T=277 was the largest-ever determined T-number among icosahedral viruses. Although the T-number of the capsid lattice of Mimivirus is likely to be even larger and is suggested to be in the range of 972 ≤ T ≤ 1200, the exact value has not yet been identified. The structure of the Mimivirus particle has been determined using cryo-EM SPA without allowing the determination of the T-number of the capsid lattice due to the low resolution resulting from the large size and the asymmetry of the virion (Xiao et al., 2005, Xiao et al., 2009, Kuznetsov et al., 2010).

## Results

### Identification of structural proteins in the MelV particle

We analyzed the protein components of purified Melbournevirus (MelV) particles by SDS-PAGE. The MelV virions contain at least 10 major proteins in abundance and many minor proteins (Fig. 1A). The most intense band appears around 50 kDa and is thought to correspond to the MCP (MW 52.4 kDa), which is conserved across many NCLDVs (black arrow in Fig. 1A).

**Fig. 1.**
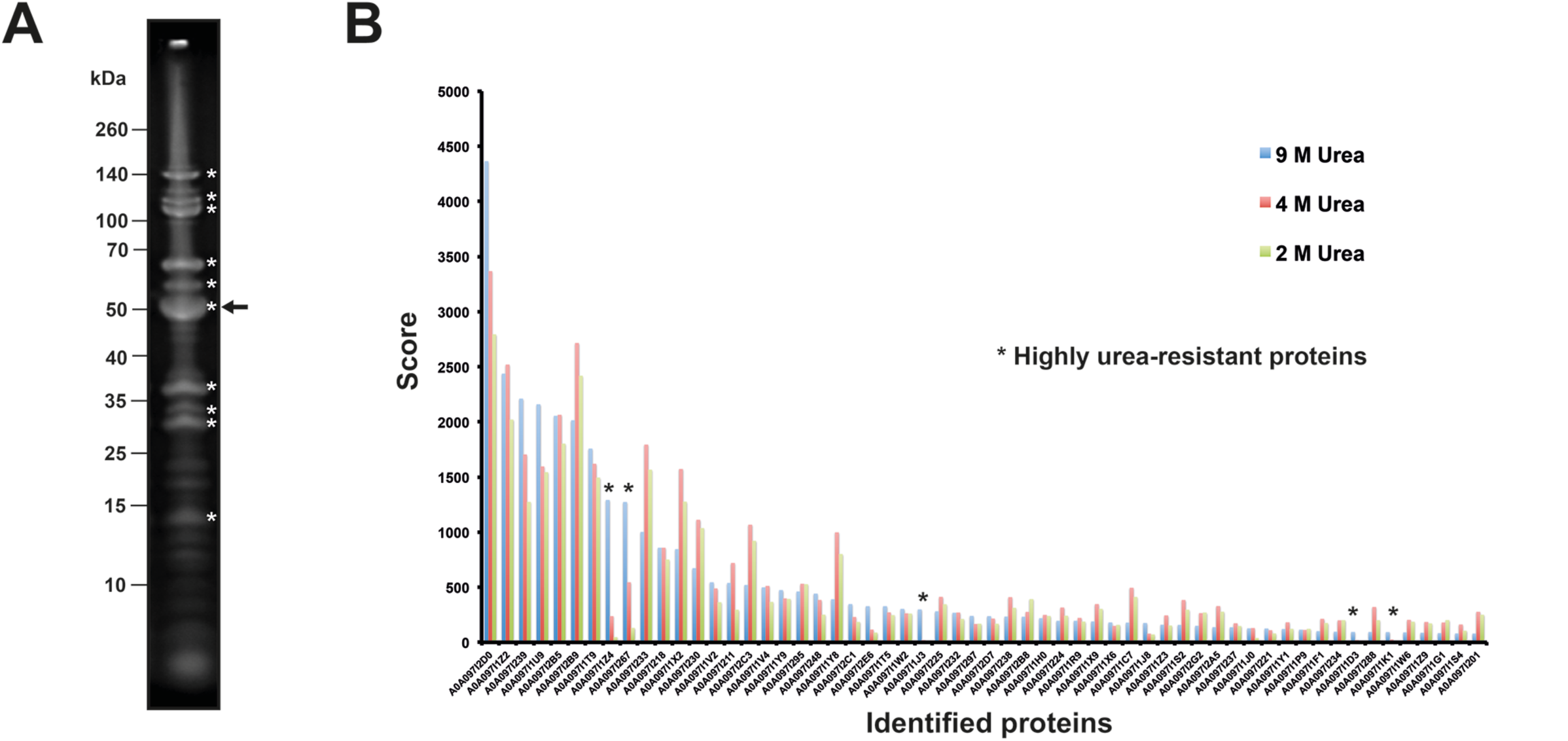
Proteins composing the MelV virions. A) SDS-PAGE analysis of purified virions. The arrow indicates the expected band of the conserved MCP. White asterisks indicate 10 abundant major proteins. B) The SEQUEST scores of the 58 identified structural proteins after treatment with 2, 4 or 9 M urea buffers. The accession codes for the identified proteins correspond to entries in the Table S1. Five proteins had reduced SEQUEST scores under 4 and 2 M urea conditions and were considered highly urea-resistant (asterisks).

The purified virus particles were also analyzed by tandem mass spectrometry (MS)-based proteomics analysis after 9 M urea treatment (Table S1) and 58 MelV-encoded proteins were identified with high reliability (SEQUEST score > 80) including three histone-like or histone-related proteins. In addition, we identified several putative viral enzymes: a cysteine peptidase, helicases, AAA-family ATPases, thioredoxins, putative serine/threonine protein kinases, an uracil-DNA glycosylase, a putative glycosyltransferase, a mannosyltransferase, an amine oxidase, a putative ribonuclease, a sulfhydryl oxidase, 31 functionally uncharacterized proteins, and 6 putative membrane proteins. Proteins probably deriving from the acanthamoeba host such as actin (Accession number (AN): L8GT00, P02578) and a few stress-induced proteins were also identified with relatively low scores. Further characterization of the structural proteins was conducted by treating the sample with varying urea concentrations. As a result, we identify five proteins that can be considered highly urea-resistant (Fig. 1B), including the conserved MCP (MEL_305, AN: A0A097I267, Score 1273). The capsid was robust enough to partially withstand a 4 M urea and thus the incompletely dispersed lattice of MCP trimers might simultaneously have pulled down four other proteins during the centrifugation of the sample washing (Asterisks in Fig. 1B, MEL_236 - AN: A0A097I1Z4, MEL_089 - AN: A0A097I1J3, MEL_025 - AN: A0A097I1D3, MEL_096 - AN: A0A097I1K1). It is worth noticing that MEL_236 (AN: A0A097I1Z4), a 16 kDa predicted protein, seems to be as abundant as the MEL_305 MCP (score 1292).

### Cryo-EM and single particle reconstruction

The size of the icosahedral MelV particle is ~230 nm in diameter (Fig. 2A). An approximately spherical double layer, assumed to be an inner membrane, could be observed beneath the icosahedral capsid (white arrows in Fig. 2B). The layer seems more diffuse around the 5-fold axes (Fig. 2C). Almost all particles show a dense interior, which indicates that most of them are filled with the viral genome and proteins. All particles clearly display a dense interior spot, approximately 30 nm in diameter. The shape of the large dense body (LDB) does not seem to be spherical and the location within the particle varies, but it is consistently observed (black arrows in Fig. 2A and a white circle in Fig. 2B).

**Fig. 2.**
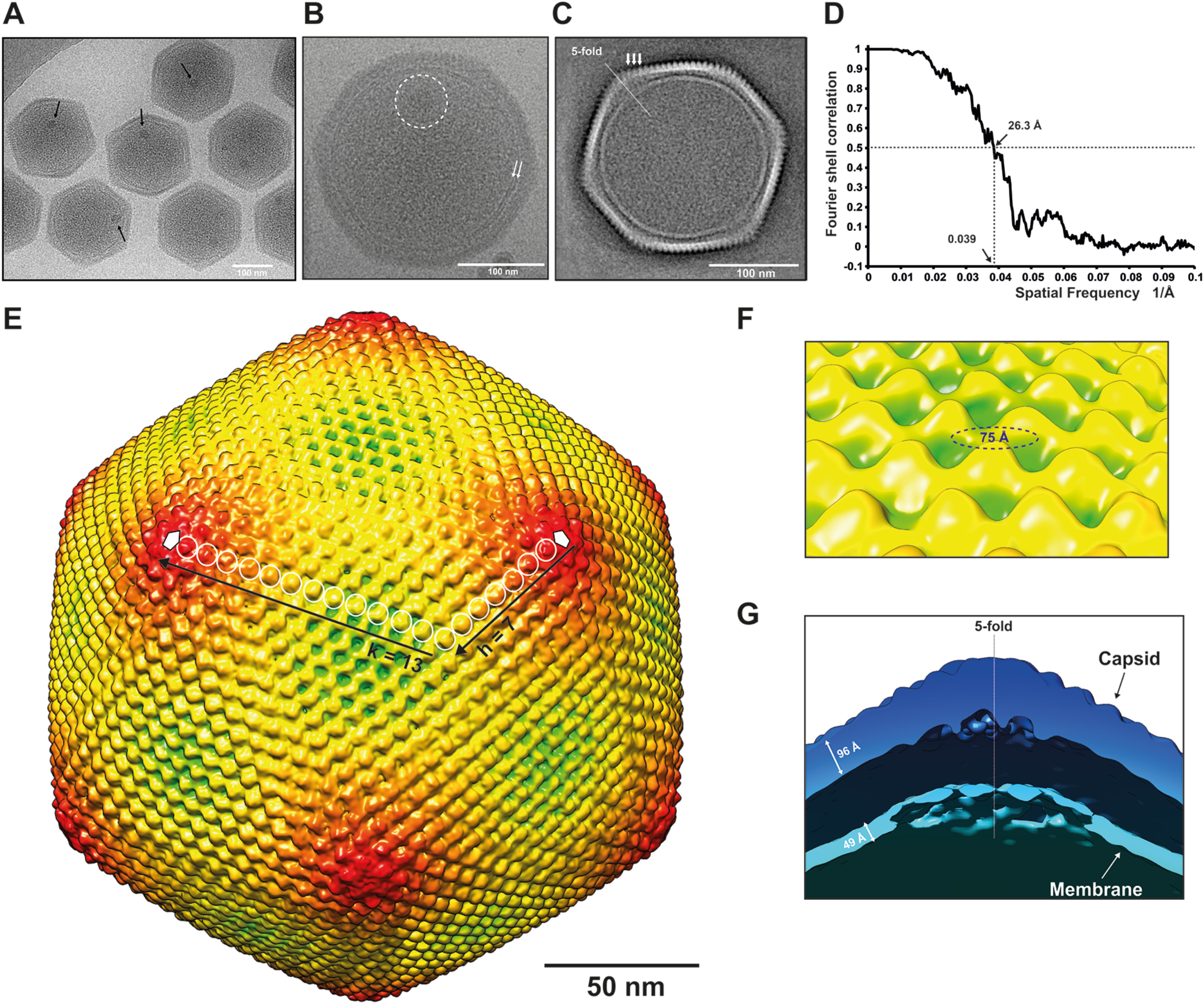
Images of cryo-frozen MelV particles and single particle 3D analysis of the particle. A, B) Raw and C) 2D class-averaged images of the MelV. Black arrows in A) and white dotted circle in B) indicate LDBs. White arrows in B) indicate the double-layered membrane. White arrows in C) indicate periodically spaced capsomers in the capsid lattice. D) Resolution plot of the single particle 3D reconstruction calculated by the FSC method with a 0.5 cutoff. E) The T=309 capsid lattice of the MelV. Surface plot of the reconstructed 3D structure coloured according to the distance from the centre of the virus particle (green < 1050 Å, yellow < 1070 Å, red < 1200 Å. The T-number was determined by the h and k numbers of the capsid lattice (back arrows and white circles). Each protrusion represents a capsomer. T=309 can be obtained by either h=7 and k=13 or h=13 and k=7. F) Close-up view of the surface protrusions that are expected to be psudo-hexagonal capsomers. The capsid structure was rendered at an isodensity contour level of 2.0 *σ* using a cryo-EM model at the resolution of 26.3 Å with an applied inner mask (E and F). G) A cross-section of the 3D density perpendicular to a 5-fold axis. The capsid and membrane structures were rendered at an isodensity contour level of 1.0 *σ* using a cryo-EM model at the resolution of 35.0 Å without applying any inner mask.

Icosahedral symmetry was imposed during classification as well as averaging of the particles for 3D reconstruction. The spherical layer is clearly observed beneath the icosahedral capsid in re-projections from the 3D reconstruction, but the density was weaker and the double layer structure smeared out around the 5-fold vertices (Fig. 2C). The LDB and other interior features were also canceled out in the symmetrized re-projections. However, the re-projection clearly shows a periodical lattice structure of the capsid (white arrows in Fig. 2C). In total, 7,005 cryo-EM projection images of virus particles were used for the 3D reconstruction and a resolution of 26.3 Å was acquired (the resolution was estimated on the basis of the 0.5 Fourier shell correlation (FSC) cutoff (Fig. 2D). The distance between the two farthest 5-fold vertices of the capsid is 232 nm. The capsid surface is covered with ordered protrusions (Fig. 2E). Similarly ordered protrusions have previously been reported in the structures of other related NCLDVs at intermediate resolution, and assigned as the pseudo-hexagonal capsomers of the conserved trimeric MCP (Nandhagopal et al., 2002, Yan et al., 2005). The ordered protrusions of the MelV particle are approximately 75 Å in diameter (Fig. 2F), which is in the range of the capsomer sizes of other NCLDVs (Nandhagopal et al., 2002). The typical thicknesses of the capsid and the membrane are 96 Å and 49 Å, respectively (Fig. 2G). The 49 Å-thickness of the double layers underneath the capsid is consistent with a typical thickness of lipid membrane bilayers. The internal membrane is thicker and appears to bulge out near the 5-fold axis (Fig. 2G).

### Capsid lattice and the T-number

The triangulation number T describes the number of identical structural units S = 60T with equivalent (or quasi-equivalent for T>1) environment in the asymmetric unit of an icosahedral capsid, as first described by Caspar and Klug (Caspar and Klug, 1962). The allowed T-numbers are defined by the formula T = Pf^2^ where P = h^2^ + hk + k^2^ (the integers h and k define the number of lattice points between neighboring vertices) and for any integer of f. Casper and Klug categorized the T-numbers into three different classes: P = 1 (T = 1, 4, 9, 16…), P = 3 (T = 3, 12, 27, 48…) and the skew class with P ≥ 7 (T = 7, 13, 19, 21…). They also correctly predicted that structural units can cluster into hexameric and pentameric groups, giving rise to M distinct morphological units or capsomers, with M being the sum of 10 x (T-1) hexamers and exactly 12 pentamers. At the given resolution of the reconstruction, the hexameric capsomers of the MelV particle appear as ordered protrusions (Fig. 2F). Counting the number of capsomers in two neighboring 5-fold vertices (Fig. 2E), we conclude that T = 309 for the MelV capsid lattice. Since 309 is a prime number, f must be 1 and therefore P ≥ 7. Hence, the lattice belongs to a skew class. That gives two options for h and k; either h = 7 and k = 13 or h = 13 and k = 7.

### Pentasymmetrons and trisymmetrons model

Wrigley first observed that collapsed capsids of large viruses tend to disintegrate into pentagonal and triangular units where the triangular units do not necessarily correspond to the triangular facets of the approximately icosahedral capsid (Wrigley, 1969). He referred to these units as pentasymmetrons and trisymmetrons, and the full capsid is built up by 12 pentasymmetrons centered at the 5-fold axes, and by 20 trisymmetrons centered at the 3-fold axes. This structural organization has since been observed in other NCLDVs (Yan et al., 2000, Yan et al., 2005). A theoretical analysis of the possible pentasymmetron-trisymmetron models of large NCLDVs was previously published (Simpson et al., 2003). According to the theory, the size of the pentasymmetrons and trisymmetrons correlates with the h and k values used for T-number determination. Since the MelV lattice belongs to the skew class it may adopt two different sets of h and k values corresponding to distinct pentasymmetron-trisymmetron models (Fig. S1). In the case (h, k) = (7, 13), the lattice consists of pentasymmetrons with 4 capsomer long edges, and trisymmetrons with 16 capsomer long edges (model in Fig. S1, left), giving 30 and 136 capsomers in each pentasymmetron and trisymmetron, respectively (Table 1). Alternatively, if (h, k) = (13, 7), the pentasymmetrons have 7 capsomer long edges, and the trisymmetrons have 13 capsomer long edges (model in Fig. S1, right). In either case, the total number of pseudo-hexagonal capsomers is 3,080 (Table 1), which in turn are assembled by 9,240 MCPs in the entire capsid lattice. Information of the orientation of the pseudo-hexagonal capsomers is required for deducing which pentasymmetron-trisymmetron model is correct for the MelV particle. Although it is difficult to determine the orientations of the capsomers at the current resolution, the model that is built with h=7 and k=13 is structurally and evolutionary more likely (see discussion section for details), hence we will only consider this model hereafter.

**Table 1.**
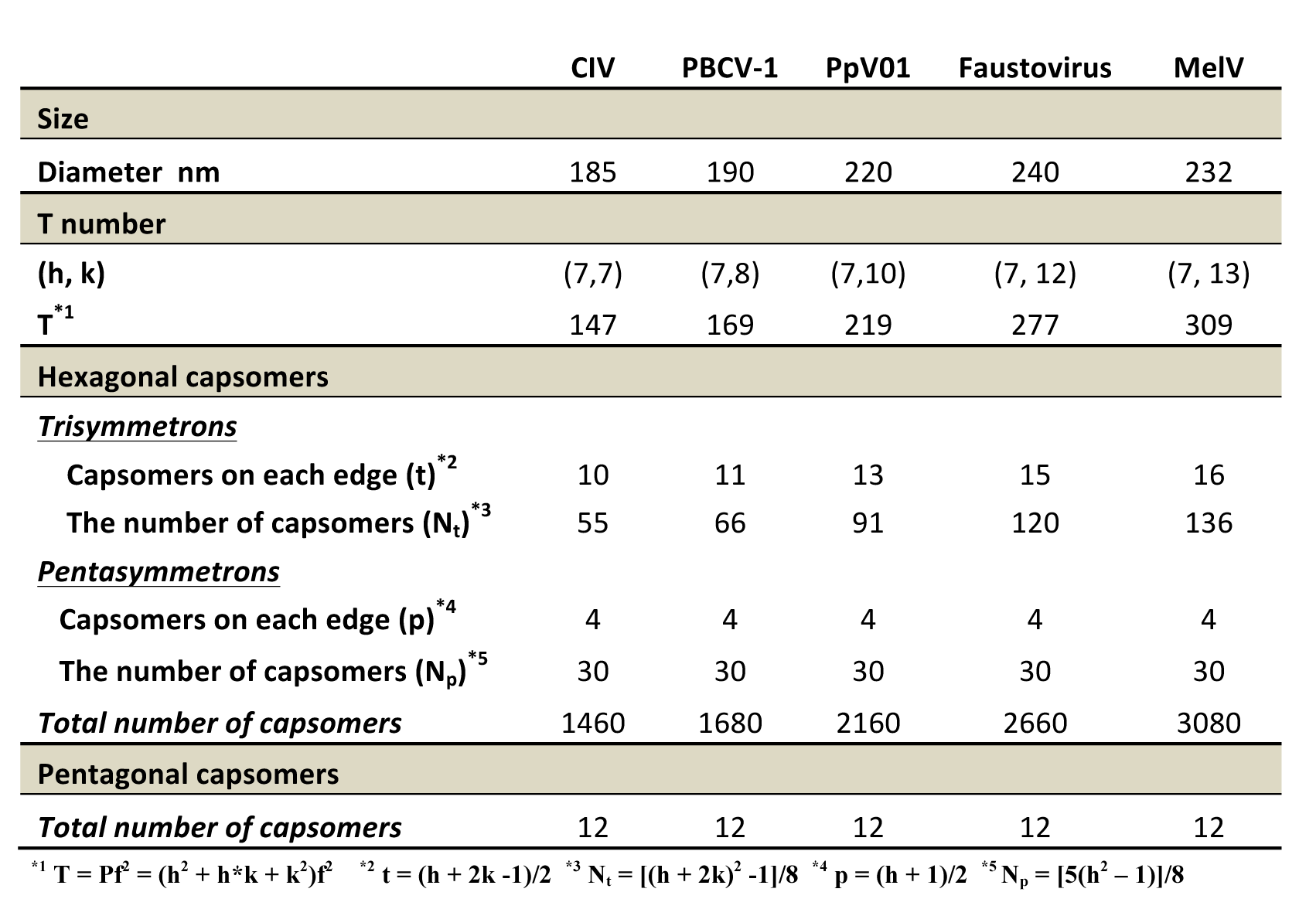
Pentasymmetron and trisymmetron organization in five giant viruses

### Other features of the capsid

The capsid structure of the MelV particle has several unique features, which cannot be explained merely by the simple quasi-equivalence model of hexagonal trimers of the MCP. The ordered capsomers on the face of the capsid, near the 3-fold axis, are oriented straight up (Fig. 3B), whereas the capsomers near the 5-fold axis seem to have varying orientations (Fig. 3A). The capsid also seems denser (red capsomers in Fig. 3C) along the edges of icosahedral surface, near the 2-fold axis. These capsomers do not show large empty spaces between the protrusions at lower isodensity levels, unlike those seen closer to the 3- and 5-fold axes (Fig. 3A-C). A superimposition of the cross-sections of the capsomers near the 2-fold and 3-fold axes shows that the additional denser regions exist along the edges of the 2-fold axes (orange densities in Fig. 3D). The denser areas near the 2-fold axes are located at the intersecting boundary between two trisymmetrons (red-edged hexagons in Fig. 3E). The maximum likelihood-based 2D class-averaged image shows the additional anchoring substructures between the capsid and the bulged out membrane underneath the pentasymmetrons near the 5-fold axes (Fig. 4).

**Fig. 3.**
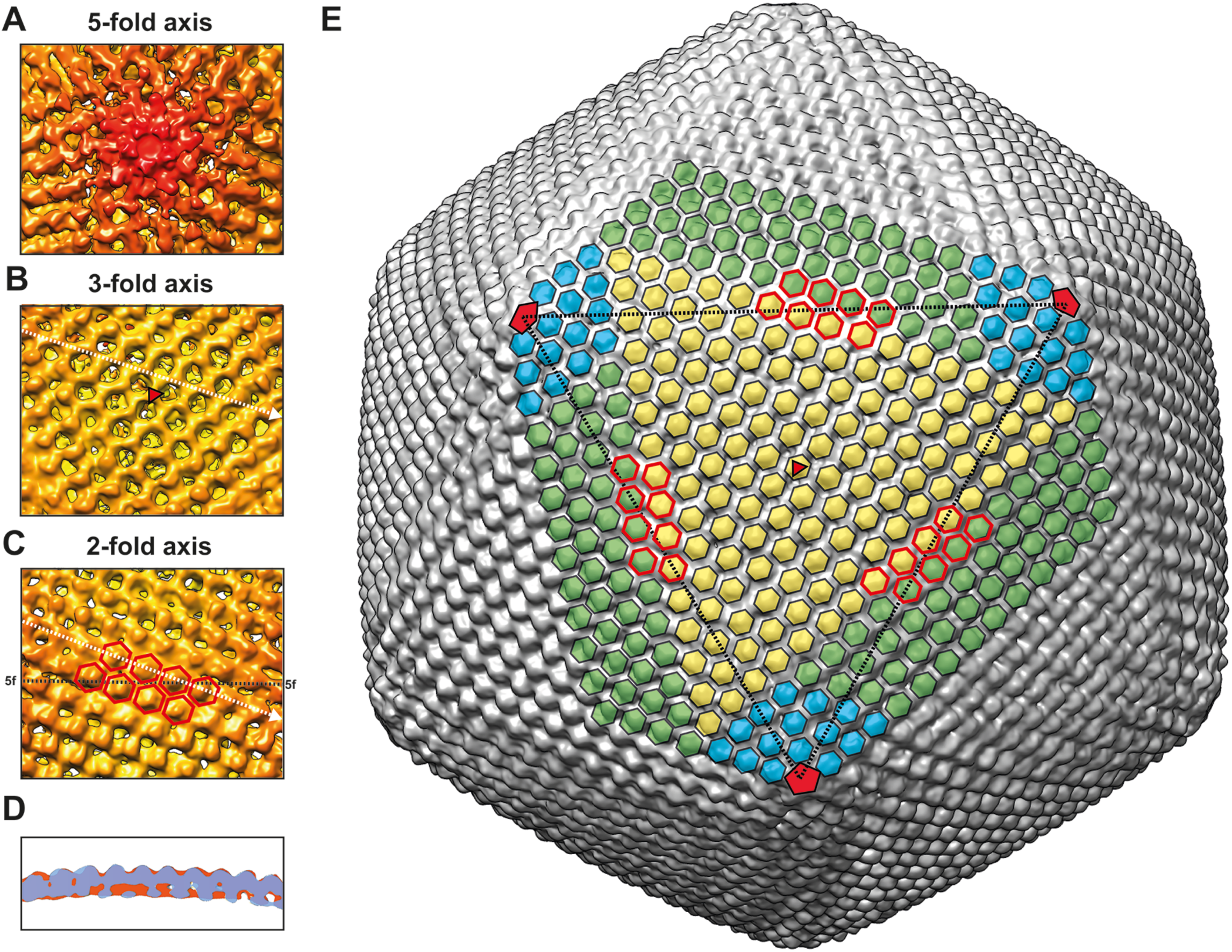
Minor capsid proteins of the MelV particle. Close-up views from A) a 5-fold, B) a 3-fold and C) a 2-fold axis of the capsid. A-C) The close-up views were rendered at a higher contour level than in Fig. 2E (an isodensity contour level of 2.5σ) to emphasize minor features. D) A superposition of cross-sections of capsomers at the 2-fold (surface in orange) and 3-fold (surface in light blue) axes. The cross-sections were generated through the white dotted lines in B and C. E) Locations of the dense capsomers, pentasymmetron, and trisymmetrons. The red-edged hexagons are the capsomers near the 2-fold axis and correspond to those in C). The black-dotted lines indicate the lines between the adjacent 5-fold axes. A red triangle and pentagons indicate the 3-fold and the 5-fold axes.

**Fig. 4.**
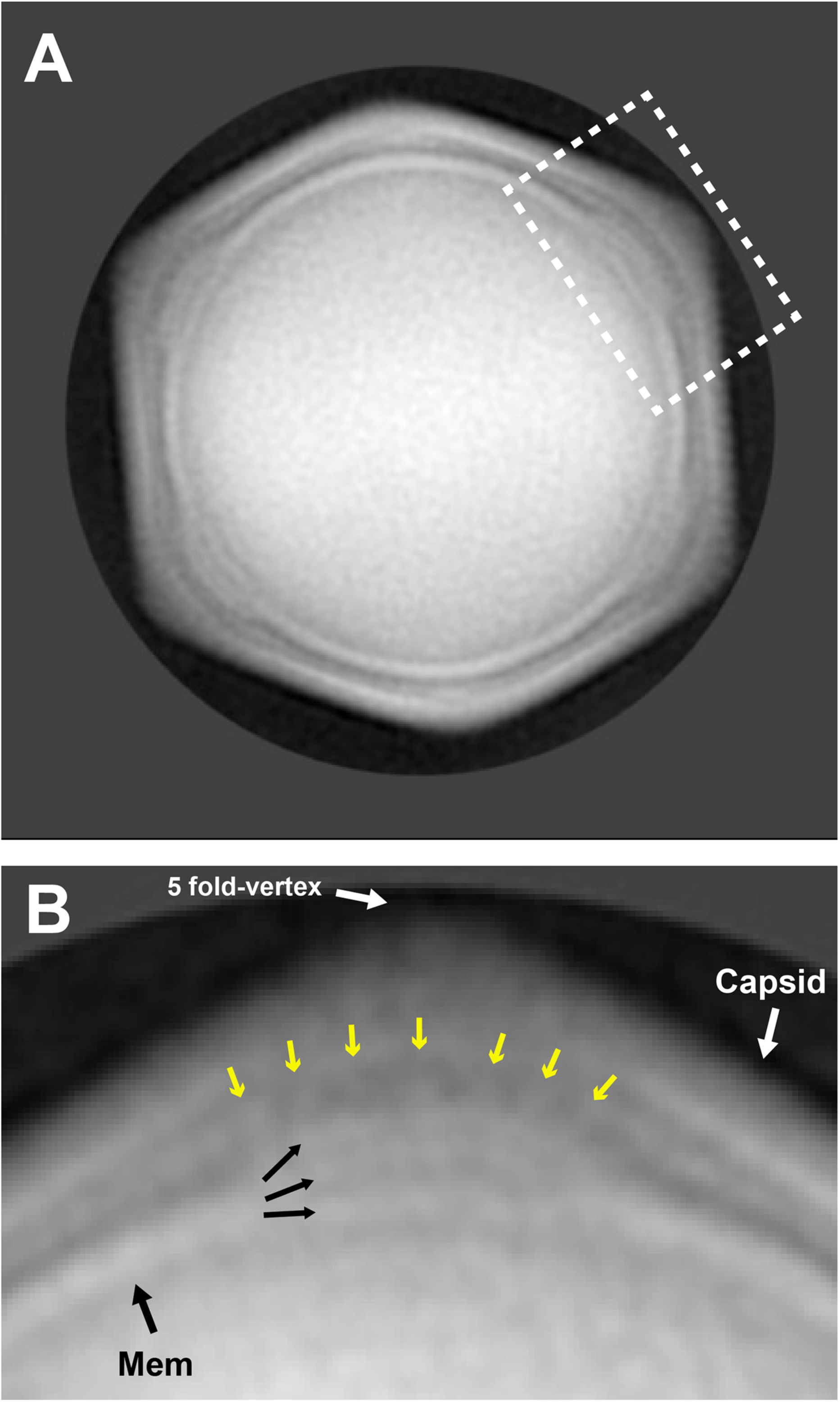
A 2D class-averaged projection of the MelV particle by maximum likelihood-based 2D classification. A) 2D class-averaged projection and B) close-up view of the 5-fold vertex corresponding to the white dotted area in A). Black arrows indicate the membranes (Mem) and the arrangement in the 5-fold vertex. Yellow arrows indicate the possible anchoring between the membrane and capsid near 5-fold vertex.

### Distribution of LDBs analyzed by electron cryo-tomography

3D models (subtomograms) for 147 complete virus particles were obtained by electron cryo-tomography (cryo-ET). In contrast to the 3D model obtained by SPA (Fig. 2G), the resolution in cryo-ET was too low to distinguish the capsid and the double-layered membrane (white isosurface in Fig. 5A). The LDB appears as a compact object with high density in an off-centered position close to the capsid/membrane (green isosurface in Fig. 5A). Assuming the weak phase approximation (Vulovic et al., 2014), the image intensity is locally proportional to the object density. The density of the LDB is estimated to be around 1.60 g/cm^3^ in comparison to the average intensity of the vitreous ice (density 0.92 g/cm^3^) and the peak intensity of the capsid (typical protein density 1.36 g/cm^3^) (Spahn et al., 2000, Perlman et al., 1982). The rest of the interior regions have a density in the range of 1.10–1.30 g/cm^3^ (Fig. 5B). Multiple 3D subtomograms of MelV particles were aligned to one unique vertex that is the closest to the LDB, and these aligned subtomograms were averaged (Fig. 5C). The LDB is most commonly located near the capsid/membrane (Fig. 5C). The 3D probability distribution of the LDB is calculated from the coordinates of the vertices and the LDB (Figs. 5D, 5E, Movie S1). The majority of the LDBs are located 400 Å from the closest vertex and 166 Å under the membrane (Fig. 5E). The LDBs are rarely located exactly on the 5-fold axes (Fig. 5D). A small number of LDBs are located near the center of the particles (Fig. 5D).

**Fig. 5.**
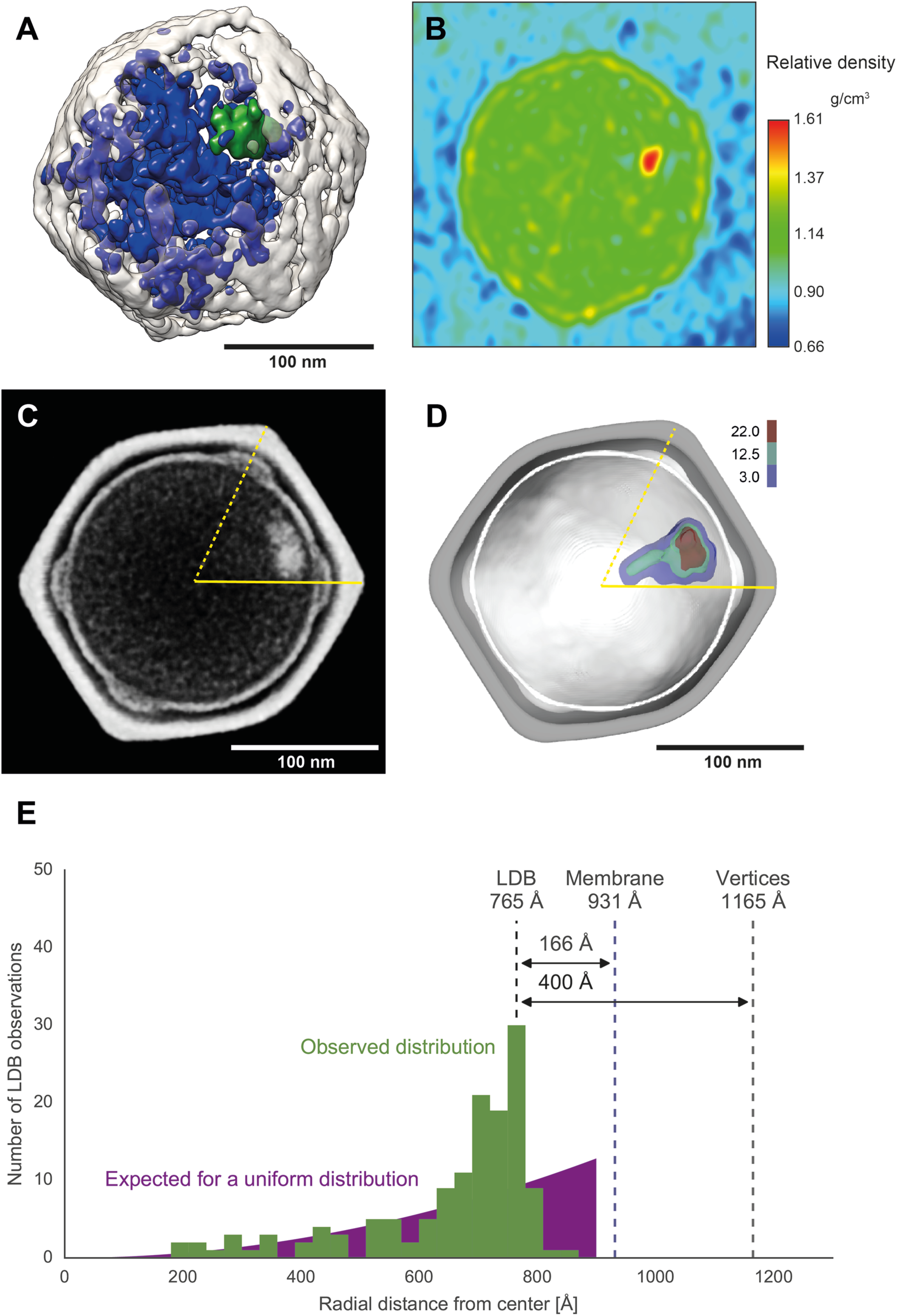
Cryo-ET 3D reconstruction of the MelV particle, and the intensity and the position of the LDB. A) One of the 3D tomographic reconstructions. The outer capsid and the membrane, the LDB, and the other interior fragments are shown with white, green, and blue isosurfaces, respectively. B) The relative image intensity in a cross-section of the virion in A), which corresponds to the relative density of the specimen in a weak phase approximation. C) A cross-section of a subtomogram-averaged model of the MelV particle. D) Spatial distribution of the LDB. The grey and white surfaces indicate the capsid and membrane of the SPA model, respectively. The probability of the spatial distribution (per Å^3^) of the LDB in the MelV particle is shown in a color gradient (3.0 – 22.0 x 10^9^/Å^3^). C, D) The 5-fold axes are indicated with yellow lines (The closest 5-fold axis of the LDB is indicated with a solid yellow line). E) Distribution of the radial distance of the LDB from the closest 5-fold vertex and the membrane (Green histogram). The purple curve shows the probability of finding the LDB, assuming a uniform distribution in the MelV particle.

## Discussion

The resolution of the MelV cryo-EM model as a fraction of Nyquist frequency is 0.25 (magnification scale: 3.31 Å/pixel, achieved resolution: 26.3 Å), which is lower than those of other reported cryo-EM models using a CCD (Chen et al., 2008). Particle heterogeneity, and weak signals coming from the thick sample may have limited the resolution here. Furthermore, since the MelV particle is large, only a handful of MelV virions fit within the field of view at high magnification, making data collection more time consuming than usual. In this study, this restriction limited the number of recorded images and hampered our efforts in improving resolution.

The MelV structure determined here is the first and the largest interpretable 3D structure of a member of the *Marseilleviridae* family (Fig. 2). The diameter of the MelV particle (232 nm) is 12 nm larger than that of PpV01 (220 nm) (Yan et al., 2005), corresponding to a 1.2 times larger volume, while the genome size of MelV (369,360 bp: 369-kbp) is significantly smaller than that of PpV01 (485-kbp) (Doutre et al., 2014, Monier et al., 2008). Why does MelV require such a large particle despite harboring a significantly smaller DNA genome in comparison to other NCLDVs? The *Mar seilleviridae* viruses uniquely encode histone-like proteins (Doutre et al., 2014, Thomas et al., 2011, Aherfi et al., 2014a), and we identified three histone-like or histone-related proteins in the purified particle (Fig. 1, Table S1). *Mar seilleviridae* viruses may have a specific genome-packaging mechanism with a histone-like superstructure of their dsDNA genome.

According to theory, the capsid lattice of the MelV particle may belong to two possible lattices (Fig. S1). To conclusively distinguish between these two cases, a higher resolution model has to be produced, to reveal the specific details of the inter-capsomer connectivity network (Simpson et al., 2003). Other NCLDVs have pentasymmetrons composed of 4 capsomer long edges (Table 1). The evolution towards larger capsids merely requires increasing the size of the trisymmetrons (Fig. 6). One of the MelV models (Fig. S1, left model) is consistent with this pattern. The other MelV model (Fig. S1, right model) would require concerted adaptations of the pentasymmetron and the trisymmetron sizes as well as of the interfaces between them. Thus, the model with small pentasymmetrons (Fig. S1, left model) is evolutionary and structurally more likely, by parsimony.

**Fig. 6.**
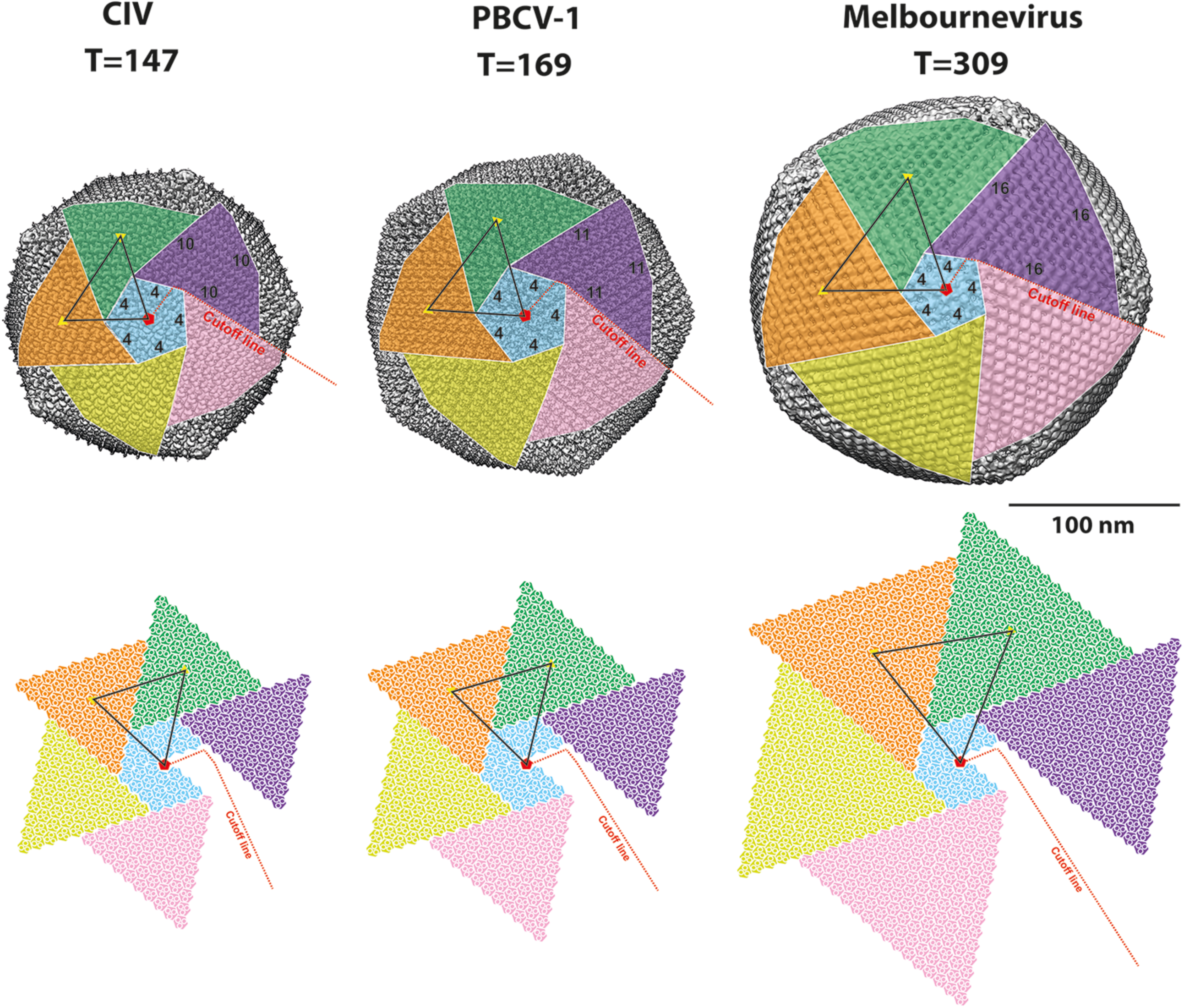
Trisymmetron and pentasymmetron organization of CIV (EMDB Accession code: 1580), PBCV-1 (EMDB Accession code: 5378) and MelV particles. The number of capsomers on each edge are shown in the figure. Yellow triangles indicate 3-fold axes and a red pentagon a 5-fold axis. Three white circles in every hexagonal capsomers (the bottom expansion plains) indicate the trimeric MCPs that compose the capsomer.

The 232 nm-sized capsid of the MelV particle is the largest among known *Marseilleviridae* viruses. In comparison with Cannes 8 virus, another member of the *Marseilleviridae,* is less than 200 nm in diameter (Aherfi et al., 2014b). Further studies of other smaller *Marseilleviridae* virions are required to clarify the relationship between the capsid size and the sizes of pentasymmetrons and trisymmetrons. The Mimivirus particle is expected to form a capsid with a much larger T-number lattice than any other known viruses (Xiao et al., 2005, Xiao et al., 2009). However, the sizes of its pentasymmetrons and trisymmetrons are still unclear. Further structural analysis of the Mimivirus particle at a higher resolution may elucidate the evolutional and structural strategies leading to such giant icosahedral capsid lattices. Minor capsid proteins are important in several other NCLDVs. PBCV-1 and PpV01 particles have 60 bean-like protrusions on certain capsomers, which likely play a role in cell receptor recognition (Yan et al., 2005, Yan et al., 2000). Chilo iridescent virus (CIV) and PBCV-1 particles have finger-like proteins on the capsomers, and CIV and Faustovirus particles also have zipper and anchor proteins, which may function as “cement” or “glue” to connect capsomers (Yan et al., 2009, Klose et al., 2016). Such minor proteins have also been described in several other dsDNA viruses, such as P30 and P16 in bacteriophage PRD1 (Abrescia et al., 2004, San Martin et al., 2002), P1 (expected anchor protein) in PM2 (Huiskonen et al., 2004), and protein IIIa (or C-termini of protein IX) and pVIII in adenovirus (Nemerow et al., 2012, Liu et al., 2010). The additional denser regions at the 2-fold axes in the MelV capsid (Fig. 3B-D) might be composed by such minor protein(s). The denser zones are located at the intersection of the boundary between two adjacent trisymmetrons (red-edged hexagons in Fig. 3E), indicating that the icosahedral facets are jointed like door hinges to adopt to the large curvature at the edge between icosahedral facets. The large disparity in size between the trisymmetrons and the pentasymmetrons may induce additional structural stress and require a mechanism for connecting them. These “hinge” proteins are probably related to the reported “cement” and “glue” proteins of other dsDNA viruses. The “hinge” proteins are not directly assigned by the proteomics analysis, however one or several of the urea-resistant proteins that were found to be associated with the MCPs could serve this purpose (Fig. 1B, Table S1). The orientations of the capsomers are distinct at the 5-fold axis (Fig. 3A). This may be the result of the structural diversity of the capsomers at different positions in the quasi-equivalent capsid lattice (Caspar and Klug, 1962). It has been disputed whether the *Marseilleviridae* viruses possess fibers on the surface or not (Boyer et al., 2009, Doutre et al., 2014). In our study, no such fibrous structures were observed on the MelV capsid neither in the raw cryo-EM images nor in the reconstructed icosahedrally-averaged 3D model (Fig. 2A, 2E).

Not much is known about the capsid assembly process of large dsDNA viruses. The current understanding is that the basic structural units are formed as pentasymmetrons and trisymmetrons. It has been demonstrated that these units are key assembly intermediates in the formation of capsids (Wrigley, 1969, Yan et al., 2005, Burnett, 1985). An icosahedral vertex of the Mimivirus seems to be initially generated during the capsid assembly from the periphery of the virus factory (Mutsafi et al., 2013). This indicates that an icosahedral vertex of the MelV particle may be initially built up by five trisymmetrons that are connected by “hinge” protein(s) and a pentasymmetron. The NCLDVs have been reported to encapsulate interior liposome-like vesicles that are asymmetric and flexible (Yan et al., 2000, Yan et al., 2005). In contrast, the virions of MelV had a more spherical and seemingly more rigid membrane structure (Fig. 2B, 2C). Six membrane proteins were identified in the MS-based proteomics analysis (Table S1) and some of them are probably incorporated into the interior membrane bilayer. The MelV membrane is less well-defined at the 5-fold axes and seemed to bulge out (Fig. 2G, 4B and 5C). A high degree of integral membrane proteins or anchor proteins of the capsid are likely embedded at these sites, thereby disrupting the spherical membrane structure. The anchoring between the membrane and the 5-fold vertices can be seen in the 2D class-averaged projection (Fig. 4). The CIV particle possesses a similar membrane feature and the anchor proteins tether the lipid membrane to the pentametric 5-fold vertices (Yan et al., 2009). The membrane in turn may be anchored to the 5-fold vertex aided by membrane proteins to further guide the complete assembly of the particle. However, many questions remain regarding the accurate MelV particle assembly mechanism. Additional proteins are probably needed to guide the capsid assembly of these large virions such as the putative MCP-interacting protein(s) (MEL236, MEL089, MEL025, MEL096) (Fig. 1B, Table S1).

No similar interior LDB-like complex has been reported in the structures of other NCLDVs (Yan et al., 2000, Yan et al., 2005, Reteno et al., 2015, Xiao et al., 2009). It is astonishing that a large 30 nm object, similar in size to a small virus or to a ribosome, is incorporated into a virus particle (Figs. 2A, 2B and 4). Several gigantic amoeba viruses episodically form dot-like or interior fibrillar components (Philippe et al., 2013, Legendre et al., 2014), but in MelV every single imaged virus particle had such a component. The position of the LDB was not fixed in the virion, however rarely located exactly on the 5-fold axis or near the center of the particle, and seems to be loosely restrained to the proximity of the capsid/membrane (Fig. 5D). This tethering of the LDB may be the result of an interaction between the LDB and the membrane. Expected membrane proteins may also impede the precise positioning of the LDBs below the 5-fold vertices (Fig. 4, Fig. 5D). The LDB is estimated to be around 1.2 times denser than the capsid layer of the virus (Fig. 5B). It is difficult to pack proteins into the LDB in a manner that makes them denser than the capsid lattice. A previous study of bulk cross-linked chromatin fragments (Schwartz et al., 2005) indicated a density of 1.39–1.42 g/cm^3^, while the density of cross-linked proteins is ≤1.25 g/cm^3^ and the density of free DNA is ≤1.69 g/cm^3^. The presence of several histone-like structural proteins encourages the hypothesis that the LDB consists of a nucleoprotein complex. It is however not feasible to pack the entire 369 kbp genome into such a small volume, but a part of the genome may be more densely packed or bound to transcription factors or a helicase of the MelV. The density of a eukaryotic ribosome is approximately 1.60 g/cm^3^ in free form (Vournakis and Rich, 1971) and the density of the LDB agree on that of the ribosome, although Mollivirus is the only known giant amoebal virus that incorporates host ribosomal proteins (Legendre et al., 2015). The eukaryotic nucleus forms a dense nucleolus, where it primarily serves as a site of ribosomal synthesis and assembly (Thiry and Lafontaine, 2005). Our proteomics experiments show that MelV possesses a conserved putative ribonuclease III (A0A097I1Y1 in Table S1). Bacterial and eukaryotic RNase III is required for RNA processing localizing to the bacterial nucleoid and eukaryotic nucleolus (Malagon, 2013, Wu et al., 2000). The LDB may include enzymes, transcriptions factors, and parts of the viral genome to initialize the generation of new virus particles.

## Materials and Methods

### Virus propagation and purification

MelV was propagated in *Acanthamoeba castellanii* cells cultured in PPYG medium (2.0% w/v proteose peptone, 0.1% w/v yeast extract, 4 mM MgSO_4_, 0.4 mM CaCl_2_, 0.05 mM Fe(NH_4_)_2_(SO4)_2_, 2.5 mM Na_2_HPO_4_, 2.5 mM KH_2_PO_4_, and 100 mM sucrose, pH 6.5). The protocol used to purify particles was described previously (Doutre et al., 2014). In short, the infected culture fluid (ICF) was collected and centrifuged for 10 min at 500g, 4 °C to remove cell debris. The supernatant was centrifuged for 35 min at 6,500g, C The pellet was suspended in 1 mL of PBS buffer. The suspended sample was loaded onto a 10–60% sucrose gradient and centrifuged for 90 min at 6,500g, 4 °C. The concentrated band was collected and dialyzed in PBS. The dialyzed sample was centrifuged for 35 min at 6,500 g, 4 °C and the pellet was suspended in PBS and used for further proteomics and cryo-EM analysis.

### MS/MS tandem spectrometry and database searches

The purified virus particles were divided into three equal volumes, centrifuged at 8,000 g for 45 min, 4 °C, and then the supernatant was removed. These pellets were suspended in 9, 4, or 2 M Urea buffer and the proteins were extracted. Thereafter the samples were incubated for 1 h at room temperature, sonicated, and then centrifuged at 16,000 g, 4 °C for 10 min. The supernatants were collected, reduced, alkylated and digested by Trypsin (Roche Applied Science) overnight as described elsewhere (http://www.scilifelab.se/facilities/bioanalytical-proteomics/). The digested samples were purified on a Pierce C18-column (Thermo Scientific), dried and resolved in 0.1% formic acid. The resulting peptides were separated in reverse-phase on a C18-column in a 15 mL 0.1% formic acid acetonitrile gradient (A: 0% and B: 99.9% acetonitrile) for 90 min, electrosprayed on-line to a Q Exactive Plus mass spectrometer (Thermo Finnigan) and sequentially analyzed by MS/MS tandem spectrometry. Database searches were performed using the SEQUEST algorithm (Eng et al., 1994), which is embedded in Proteome Discoverer 2.1 (Thermo Scientific) against proteins from *Melbournevirus* (Taxonomy ID: 1560514) and *Acanthamoeba castellanii* downloaded from the UniProt KB database (Aug 2016). The search criteria for protein identification were set to at least two matching peptides of 95% confidence level per protein. 121, 133 and 118 proteins were detected by this criterion in 9 M, 4 M and 2 M urea-treated samples, respectively. Proteins with a score over 80 according to the SEQUEST algorithm was considered with high reliability to be virions proteins and 58 out of 121 identified proteins were listed as such (Table S1).

### Cryo-EM image acquisition and single particle analysis (SPA)

Purified virus particles were applied onto a holey carbon grid (R1.2/1.3 Quantifoil Micro Tools GmbH, Germany) pretreated by glow-discharge, and plunged-frozen in liquid nitrogen-cooled ethane using Vitrobot Mark-IV (FEI Company, USA). The ice-embedded sample was observed using a field emission transmission electron microscope (FE-TEM), JEM2200FS (JEOL Ltd. Japan), at 45,317 times detector magnification (3.36 Å/pixel) with an accelerating voltage of 200 keV using a 4k × 4k charge-coupled device (F415, TVIPS GmbH, Germany) and an electron dose of ~20 e^-^/Å^2^. An omega-type energy filter was applied to obtain a zero loss electron beam (20 eV slit width). The underfocus values were approximately 1–5 μm. EMAN2 was used for reconstructing the 3D structures from the images (Tang et al., 2007). Virus particles were boxed separately using e2boxer.py (EMAN2). Contrast transfer function (CTF) calculations and amplitude corrections were performed on boxed particles by e2ctf py (EMAN2) and then manually verified by adjusting the defocus and B-factor values. Several random icosahedral initial models were calculated using class-averaged virus particle images. Initially 1,979 virus particles were used for the 3D reconstruction. The signals from the obtained images were weak, so binning was applied when calculating the first model. The pixel size after binning was 6.72 Å/pixel. Icosahedral symmetry was imposed during the 3D refinement. During the reconstruction process, outer radial masking, low-pass filtering, and FSC calculation were applied by the EMAN2 program. The calculations were conducted using a CPU cluster using 100–200 CPU cores per job. After 12 iterative steps of refinement, the first SPA model was generated, which was estimated to have a resolution of 35.0 Å using a FSC resolution cutoff of 0.5. Then, 7,005 particle images without binning (3.36 Å/pixel) and the 35.0 Å starting model were used for reconstructing the final model. After 10 iterative steps of refinement with inner and outer masking, the final SPA 3D model was generated and the reconstruction estimated to have a resolution of 26.3 Å with a FCS resolution cutoff of 0.5. The 3D volume of the final model was visualized by UCSF Chimera (Pettersen et al., 2004). A 2D class-averaged projection of the MelV particle was also generated by maximum likelihood-based 2D classification using RELION (Scheres, 2012).

### Tomographic reconstruction and data analysis

Five microliters of the virus sample was mixed with the same amount of 15 nm fiducial marker gold colloids. The specimen was loaded onto a holey-carbon grid (R1.2/1.3 Quantifoil) and plunged-frozen in liquid nitrogen-cooled ethane, and imaged with a FE-TEM, JEOL JEM2200FS, at 24,116 times detector magnification on a 4k × 4k charge-coupled device (TVIPS, F415) with an accelerating voltage of 200 kV. Specimens were tilted from 70° to -70° with 2° angular steps. Images were binned up to 12.11 Å/pixel after collecting the data. The alignment of the images and the final tomograms were calculated by the IMOD software (Kremer et al., 1996) by using the colloidal gold fiducial markers. A total of 147 subtomograms of the MelV particle were picked up by EMAN2 (Tang et al., 2007). These particles were low-pass Gaussian filtered by e2proc3d.py (EMAN2). The subsequent post-tomographic image processing was performed using EMAN1 (Tang et al., 2007, Ludtke et al., 1999). The 20 Å low-pass filtered subtomograms were aligned to the SPA model of the MelV particle. These aligned subtomograms were symmetrically rotated to place the LDB to the same asymmetric unit of the icosahedral capsid. The subtomogram-averaged model of the MelV particle was calculated by averaging the processed subtomograms. To characterize positions of individual LDBs, the coordinates of the vertices and the interior LDB were determined by the following procedure. First, the final SPR model was rotated to match the orientation of the each low-resolution subtomogram particle. Then the coordinates of the vertices of the capsid of the SPA model and the densest region within the LDB in the tomographic model were determined by manual selection using the Chimera software (Pettersen et al., 2004). The probability distribution of the LDB was calculated by the coordinates of 147 LDBs positions in the MelV particle with applying an isosurface level to these LDB positions.

## Acknowledgements

We are grateful to Chan Xiao, Department of Chemistry, University of Texas at El Paso (UTEP), for a kind discussion of determining T-number. This work was supported by the following agencies: The Swedish Research Council (to J.H., grant number: 628–20081109, 822–2010-6157, 822–2012-5260, and 828–2012-108), the Knut and Alice Wallenberg Foundation (to J.H., grant number: KAW-2011.081), the European Research Council (to J.H., grant number: ERC-291602), the Rontgen-Angstrom Cluster (to J.H., grant number: 349–2011-6488, and 2015–06107), the Swedish Foundation for International Cooperation in Research and Higher Education (STINT) (to J.H. and K.O., grant number: JA2014-5721), The European Regional Development Fund (ELIBIO CZ.02.1.01/0.0/0.0/15_003/0000447) at the European Extreme Light Infrastructure (to J.H.), KAKENHI from the Ministry of Education, Culture, Sports, Science and Technology of Japan (to N.M., grant Number: 25251009) and the Collaborative Study Program of National Institute for Physiological Sciences (to K.O., grant Number: 2016-No. 38), the CNRS, Aix-Marseille University and the French National Research Agency (to C.A., grant number: ANR-14-CE14-0023–01).

## Author contributions

JH, CA, KO, JMC, KM and MS conceived the experiment. CA and JMC isolated the virus and provided samples. KO, HMKNR, MS performed proteomics studies. KO, NM, MFH, KM and MS performed the cryo-EM studies. KO, NM, MFH, FRNCM, DSDL, CA, JMC, KM and MS analysed the data. All authors read and approved the final manuscript.

## References

Abergel, C., Legendre, M. & Claverie, J. M. 2015. The rapidly expanding universe of giant viruses: Mimivirus, Pandoravirus, Pithovirus and Mollivirus. FEMS Microbiol Rev, 39, 779-96.

Abrescia, N. G., Cockburn, J. J., Grimes, J. M., Sutton, G. C., Diprose, J. M., Butcher, S. J., Fuller, S. D., SAN MARTIN, C., Burnett, R. M., Stuart, D. I., Bamford, D. H. & Bamford, J. K. 2004. Insights into assembly from structural analysis of bacteriophage PRD1. Nature, 432, 68-74.

Aherfi, S., Boughalmi, M., Pagnier, I., Fournous, G., La Scola, B., Raoult, D. & Colson, P. 2014a. Complete genome sequence of Tunisvirus, a new member of the proposed family Marseilleviridae. Arch Virol, 159, 2349-58.

Aherfi, S., La Scola, B., Pagnier, I., Raoult, D. & Colson, P. 2014b. The expanding family Marseilleviridae. Virology, 466–467, 27-37.

Boyer, M., Yutin, N., Pagnier, I., Barrassi, L., Fournous, G., Espinosa, L., Robert, C., Azza, S., Sun, S., Rossmann, M. G., Suzan-Monti, M., LA SCOLA, B., Koonin, E. V. & Raoult, D. 2009. Giant Marseillevirus highlights the role of amoebae as a melting pot in emergence of chimeric microorganisms. Proc Natl Acad Sci USA, 106, 21848-53.

Burnett, R. M. 1985. The structure of the adenovirus capsid. II. The packing symmetry of hexon and its implications for viral architecture. J MolBiol, 185, 125-43.

Caspar, D. L. & Klug, A. 1962. Physical principles in the construction of regular viruses. Cold Spring Harb Symp Quant Biol, 27,1-24.

Chen, D. H., Jakana, J., Liu, X., Schmid, M. F. & Chiu, W. 2008. Achievable resolution from images of biological specimens acquired from a 4 k x 4 k CCD camera in a 300-kV electron cryomicroscope. J Struct Biol, 163, 45-52.

Claverie, J. M. & Abergel, C. 2013. Open questions about giant viruses. Adv Virus Res, 85, 25-56.

Claverie, J. M., Grzela, R., Lartigue, A., Bernadac, A., Nitsche, S., Vacelet, J., Ogata, H. & Abergel, C. 2009. Mimivirus and Mimiviridae: giant viruses with an increasing number of potential hosts, including corals and sponges. J Invertebr Pathol, 101,172-80.

Colson, P., Pagnier, I., Yoosuf, N., Fournous, G., La Scola, B. & Raoult, D. 2013. “Marseilleviridae”, a new family of giant viruses infecting amoebae. Arch Virol, 158, 915-20.

Dornas, F. P., Rodrigues, F. P., Boratto, P. V., Silva, L. C., Ferreira, P. C., Bonjardim, C. A., Trindade, G. S., Kroon, E. G., La Scola, B. & Abrahao, J. S. 2014. Mimivirus circulation among wild and domestic mammals, Amazon Region, Brazil. Emerg Infect Dis, 20, 469-72.

Doutre, G., Arfib, B., Rochette, P., Claverie, J. M., Bonin, P. & Abergel, C. 2015. Complete Genome Sequence of a New Member of the Marseilleviridae Recovered from the Brackish Submarine Spring in the Cassis Port-Miou Calanque, France. Genome Announc, 3.

Doutre, G., Philippe, N., Abergel, C. & Claverie, J. M. 2014. Genome analysis of the first Marseilleviridae representative from Australia indicates that most of its genes contribute to virus fitness. J Virol, 88,14340-9.

Eng, J. K., Mccormack, A. L. & Yates, J. R. 1994. An approach to correlate tandem mass spectral data of peptides with amino acid sequences in a protein database. J Am Soc Mass Spectrom, 5, 976-89.

Huiskonen, J. T., Kivela, H. M., Bamford, D. H. & Butcher, S. J. 2004. The PM2 virion has a novel organization with an internal membrane and pentameric receptor binding spikes. Nat Struct Mol Biol, 11, 850-6.

Iyer, L. M., Aravind, L. & Koonin, E. V. 2001. Common origin of four diverse families of large eukaryotic DNA viruses. J Virol, 75,11720-34.

Klose, T., Reteno, D. G., Benamar, S., Hollerbach, A., Colson, P., La Scola, B. & Rossmann, M. G. 2016. Structure of faustovirus, a large dsDNA virus. Proc Natl Acad Sci USA, 113, 6206-11.

Koonin, E. V. & Yutin, N. 2010. Origin and evolution of eukaryotic large nucleo-cytoplasmic DNA viruses. Intervirology, 53, 284-92.

Kremer, J. R., Mastronarde, D. N. & Mcintosh, J. R. 1996. Computer visualization of three-dimensional image data using IMOD. J Struct Biol, 116, 71-6.

Kuznetsov, Y. G., Xiao, C., Sun, S., Raoult, D., Rossmann, M. & Mcpherson, A. 2010. Atomic force microscopy investigation of the giant mimivirus. Virology, 404,127-37.

La Scola, B., Audic, S., Robert, C., Jungang, L., De Lamballerie, X., Drancourt, M., Birtles, R., Claverie, J. M. & Raoult, D. 2003. A giant virus in amoebae. Science, 299, 2033.

Legendre, M., Bartoli, J., Shmakova, L., Jeudy, S., Labadie, K., Adrait, A., Lescot, M., Poirot, O., Bertaux, L., Bruley, C., Coute, Y., Rivkina, E., Abergel, C. & Claverie, J. M. 2014. Thirty-thousand-year-old distant relative of giant icosahedral DNA viruses with a pandoravirus morphology. Proc Natl Acad Sci USA, 111, 4274-9.

Legendre, M., Lartigue, A., Bertaux, L., Jeudy, S., Bartoli, J., Lescot, M., Alempic, J. M., Ramus, C., Bruley, C., Labadie, K., Shmakova, L., Rivkina, E., Coute, Y., Abergel, C. & Claverie, J. M. 2015. In-depth study of Mollivirus sibericum, a new 30,000-y-old giant virus infecting Acanthamoeba. Proc Natl Acad Sci USA, 112, E5327-35.

Liu, H., Jin, L., Koh, S. B., Atanasov, I., Schein, S., Wu, L. & Zhou, Z. H. 2010. Atomic structure of human adenovirus by cryo-EM reveals interactions among protein networks. Science, 329,1038-43.

Ludtke, S. J., Baldwin, P. R. & Chiu, W. 1999. EMAN: semiautomated software for high-resolution single-particle reconstructions. J Struct Biol, 128, 82-97.

Malagon, F. 2013. RNase III is required for localization to the nucleoid of the 5’ pre-rRNA leader and for optimal induction of rRNA synthesis in E. coli. RNA, 19, 1200-7.

Monier, A., Larsen, J. B., Sandaa, R. A., Bratbak, G., Claverie, J. M. & Ogata, H. 2008. Marine mimivirus relatives are probably large algal viruses. Virol J, 5, 12.

Mutsafi, Y., Shimoni, E., Shimon, A. & Minsky, A. 2013. Membrane assembly during the infection cycle of the giant Mimivirus. PLoS Pathog, 9, e1003367.

Nandhagopal, N., Simpson, A. A., Gurnon, J. R., Yan, X., Baker, T. S., Graves, M. V., Van Etten, J. L. & Rossmann, M. G. 2002. The structure and evolution of the major capsid protein of a large, lipid-containing DNA virus. Proc Natl AcadSci USA, 99,14758-63.

Nemerow, G. R., Stewart, P. L. & Reddy, V. S. 2012. Structure of human adenovirus. CurrOpin Virol, 2,115-21.

Perlman, A. J., Stanley, F. & Samuels, H. H. 1982. Thyroid hormone nuclear receptor. Evidence for multimeric organization in chromatin. J Biol Chem, 257,930-8.

Pettersen, E. F., Goddard, T. D., Huang, C. C., Couch, G. S., Greenblatt, D. M., Meng, E. C. & Ferrin, T. E. 2004. UCSF Chimera–a visualization system for exploratory research and analysis. J Comput Chem, 25,1605-12.

Philippe, N., Legendre, M., Doutre, G., Coute, Y., Poirot, O., Lescot, M., Arslan, D., Seltzer, V., Bertaux, L., Bruley, C., Garin, J., Claverie, J. M. & Abergel, C. 2013. Pandoraviruses: amoeba viruses with genomes up to 2.5 Mb reaching that of parasitic eukaryotes. Science, 341, 281-6.

Reteno, D. G., Benamar, S., Khalil, J. B., Andreani, J., Armstrong, N., Klose, T., Rossmann, M., Colson, P., Raoult, D. & La Scola, B. 2015. Faustovirus, an asfarvirus-related new lineage of giant viruses infecting amoebae. J Virol, 89, 6585-94.

Saadi, H., Reteno, D. G., Colson, P., Aherfi, S., Minodier, P., Pagnier, I., Raoult, D. & La Scola, B. 2013. Shan virus: a new mimivirus isolated from the stool of a Tunisian patient with pneumonia. Intervirology, 56, 424-9.

San Martin, C., Huiskonen, J. T., Bamford, J. K., Butcher, S. J., Fuller, S. D., Bamford, D. H. & Burnett, R. M. 2002. Minor proteins, mobile arms and membrane-capsid interactions in the bacteriophage PRD1 capsid. Nat Struct Biol, 9, 756-63.

Santini, S., Jeudy, S., Bartoli, J., Poirot, O., Lescot, M., Abergel, C., Barbe, V., Wommack, K. E., Noordeloos, A. A., Brussaard, C. P. & Claverie, J. M. 2013. Genome of Phaeocystis globosa virus PgV-16T highlights the common ancestry of the largest known DNA viruses infecting eukaryotes. Proc Natl AcadSci USA, 110,10800-5.

Scheres, S. H. 2012. A Bayesian view on cryo-EM structure determination. J Mol Biol, 415, 406-18.

Schwartz, Y. B., Kahn, T. G. & Pirrotta, V. 2005. Characteristic low density and shear sensitivity of cross-linked chromatin containing polycomb complexes. Mol Cell Biol, 25, 432-9.

Simpson, A. A., Nandhagopal, N., Van Etten, J. L. & Rossmann, M. G. 2003. Structural analyses of Phycodnaviridae and Iridoviridae. Acta Crystallogr D Biol Crystallogr, 59, 2053-9.

Spahn, C. M., Penczek, P. A., Leith, A. & Frank, J. 2000. A method for differentiating proteins from nucleic acids in intermediate-resolution density maps: cryo-electron microscopy defines the quaternary structure of the Escherichia coli 70S ribosome. Structure, 8, 937-48.

Tang, G., Peng, L., Baldwin, P. R., Mann, D. S., Jiang, W., Rees, I. & Ludtke, S. J. 2007. EMAN2: an extensible image processing suite for electron microscopy. J Struct Biol 157,38-46.

Thiry, M. & Lafontaine, D. L. 2005. Birth of a nucleolus: the evolution of nucleolar compartments. Trends Cell Biol, 15,194-9.

Thomas, V., Bertelli, C., Collyn, F., Casson, N., Telenti, A., Goesmann, A., Croxatto, A. & Greub, G. 2011. Lausannevirus, a giant amoebal virus encoding histone doublets. Environ Microbiol, 13,1454-66.

Van Etten, J. L., Meints, R. H., Kuczmarski, D., Burbank, D. E. & Lee, K. 1982. Viruses of symbiotic Chlorella-like algae isolated from Paramecium bursaria and Hydra viridis. Proc Natl Acad Sci USA, 79, 3867-71.

Vournakis, J. & Rich, A. 1971. Size changes in eukaryotic ribosomes. Proc Natl Acad Sci USA, 68, 3021-5.

Vulovic, M., Voortman, L. M., Van Vliet, L. J. & Rieger, B. 2014. When to use the projection assumption and the weak-phase object approximation in phase contrast cryo-EM. Ultramicroscopy, 136, 61-6.

Wrigley, N. G. 1969. An electron microscope study of the structure of Sericesthis iridescent virus. J Gen Virol, 5,123-34.

Wu, H., Xu, H., Miraglia, L. J. & Crooke, S. T. 2000. Human RNase III is a 160-kDa protein involved in preribosomal RNA processing. J Biol Chem, 275, 36957-65.

Xiao, C., Chipman, P. R., Battisti, A. J., Bowman, V. D., Renesto, P., Raoult, D. & Rossmann, M. G. 2005. Cryo-electron microscopy of the giant Mimivirus. J MolBiol, 353, 493-6.

Xiao, C., Kuznetsov, Y. G., Sun, S., Hafenstein, S. L., Kostyuchenko, V. A., Chipman, P. R., Suzan-Monti, M., Raoult, D., Mcpherson, A. & Rossmann, M. G. 2009. Structural studies of the giant mimivirus. PLoS Biol, 7, e92.

Yan, X., Chipman, P. R., Castberg, T., Bratbak, G. & Baker, T. S. 2005. The marine algal virus PpV01 has an icosahedral capsid with T=219 quasisymmetry. J Virol, 79, 9236-43.

Yan, X., Olson, N. H., Van Etten, J. L., Bergoin, M., Rossmann, M. G. & Baker, T. S. 2000. Structure and assembly of large lipid-containing dsDNA viruses. Nat Struct Biol, 7,101-3.

Yan, X., Yu, Z., Zhang, P., Battisti, A. J., Holdaway, H. A., Chipman, P. R., Bajaj, C., Bergoin, M., Rossmann, M. G. & Baker, T. S. 2009. The capsid proteins of a large, icosahedral dsDNA virus. J Mol Biol, 385,1287-99.

Yutin, N. & Koonin, E. V. 2012. Hidden evolutionary complexity of Nucleo-Cytoplasmic Large DNA viruses of eukaryotes. Virol J, 9,161.

